# DeepCSO: a deep-learning network approach to predicting Cysteine S-sulphenylation sites

**DOI:** 10.1101/2020.08.12.248914

**Authors:** Xiaru Lyu, Ningning He, Zhen Chen, Yang Zou, Lei Li

## Abstract

Cysteine S-sulphenylation (CSO), as a novel post-translational modification (PTM), has emerged as a potential mechanism to regulate protein functions and affect signal networks. Because of its functional significance, several prediction approaches have been developed. Nevertheless, they are based on a limited dataset from *Homo sapiens* and there is a lack of prediction tools for the CSO sites of other species. Recently, this modification has been investigated at the proteomics scale for a few species and the number of identified CSO sites has significantly increased. Thus, it is essential to explore the characteristics of this modification across different species and construct prediction models with better performances based on the enlarged dataset. In this study, we constructed a few classifiers and fond that the long short-term memory model with the word-embedding encoding approach, dubbed LSTM_WE_, performs favorably to the traditional machine-learning models and other deep-learning models across different species, in terms of cross-validation and independent test. The area under the ROC curve values for LSTM_WE_ ranged from 0.82 to 0.85 for different organisms, which is superior to the reposted CSO predictors. Moreover, we developed the general model based on the integrated data from different species and it showed great universality and effectiveness. We provided the on-line prediction service called DeepCSO that included both species-specific and general models, which is accessible through http://www.bioinfogo.org/DeepCSO.

## 1. Introduction

Protein Cysteine S-sulphenylation (CSO) is the reversible oxidation of protein cysteinyl thiols to suphenic acids. S-sulphenylation functions as an intermediate on the path towards other redox modifications, such as disulfide formation, S-glutathionylation, and overoxidation to sulfinic and sulfonic acids [1, 2]. This modification has been reported to influence protein functions, regulate signal transduction and affect cell cycle across various species [1, 3–8]. So far, thousands of CSO sites have been identified in different species including the mammal *Homo sapiens* and the plant organism *Arabidopsis thaliana* using the chemoproteomics approach [9–13](Summarized in Table S1). Nevertheless, highly efficient detection of the CSO sites remains a major methodological issue, due to the low abundance and dynamic level of CSO-containing proteins *in vivo*. In contrast to the time-consuming and expensive experimental approaches, computational methods for predicting CSO sties have attracted considerable attention because of their convenience and efficiency.

Several computational methods have been developed for the prediction of CSO sites, mainly based on a single human dataset containing 1105 identified CSO sites [10]. They include MDD-SOH[14], iSulf-Cys[15], SOHSite[16], PRESS[17], Sulf_FSVM[18], S-SulfPred[19] Fu-SulfPred[20], SulCysSite[21], SOHPRED[22], and PredCSO[23]. They are all based on Support Vector Machine (SVM), decision tree or random forest (RF) algorithms with a variety of sequence encoding schemes. Although these algorithms have made contribution to the prediction of CSO sites, most of them are currently inaccessible. Moreover, there is a lack of prediction tools for the CSO sites of other species. With the growing number of CSO sites verified, it is essential to develop species-specific prediction models with high accuracy or even a general model.

Compared to traditional machine-learning (ML) algorithms (*e.g.,* SVM and RF) used in the prediction approaches described above, the deep-learning (DL) architecture is a promising ML algorithm. In the DL algorithm, a suitable representation of the input data can be transformed into highly abstract features through propagating the whole model. Superposition of hidden layers in neural network can increase the ability of feature extraction, resulting in more accurate interpretation of latent data pattern. Because of its excellence, DL has been recently applied in the field of Bioinformatics, such as the predictions of RNA-binding sites [24], protein structure [25], RNA modification sites[26] and various PTM sites [27–29]. In view of this, the introduction of DL algorithm into the prediction of CSO sites would be a promising move to provide reliable candidates for further experimental consideration.

In this study, we constructed a few *in silico* approaches for the prediction of the CSO sites for *H. Sapiens* and *A. thaliana*. These approaches included the RF algorithms, Convolutional Neural Network (CNN) and long short-term memory (LSTM) network. The LSTM model with word embedding, called LSTM_WE_, compared favorable to the rest approaches with AUC as 0.85 and 0.82 in human and Arabidopsis in terms of cross-validation. Moreover, LSTM_WE_ trained using the data from one species achieved outstanding performance in predicting CSO sites of other species (e.g. AUC=0.80 for the prediction of Arabidopsis CSO sites using the human model), suggesting that CSO is highly conserved. Therefore, we constructed a general CSO prediction model. These models will facilitate the discovery of new CSO sites and thus will contribute to the understanding of roles and functions of CSO in diverse cellular processes.

## 2 Materials and methods

### 2.1 Data collection and preprocessing

The experimentally identified CSO sites were derived from two different organisms including *H. Sapiens* and *A. thaliana* [9–13]. The data of the species were pre-processed and the related procedure was exemplified using the *A. thaliana* data, as listed below (Figure 1A).

**Figure 1.**
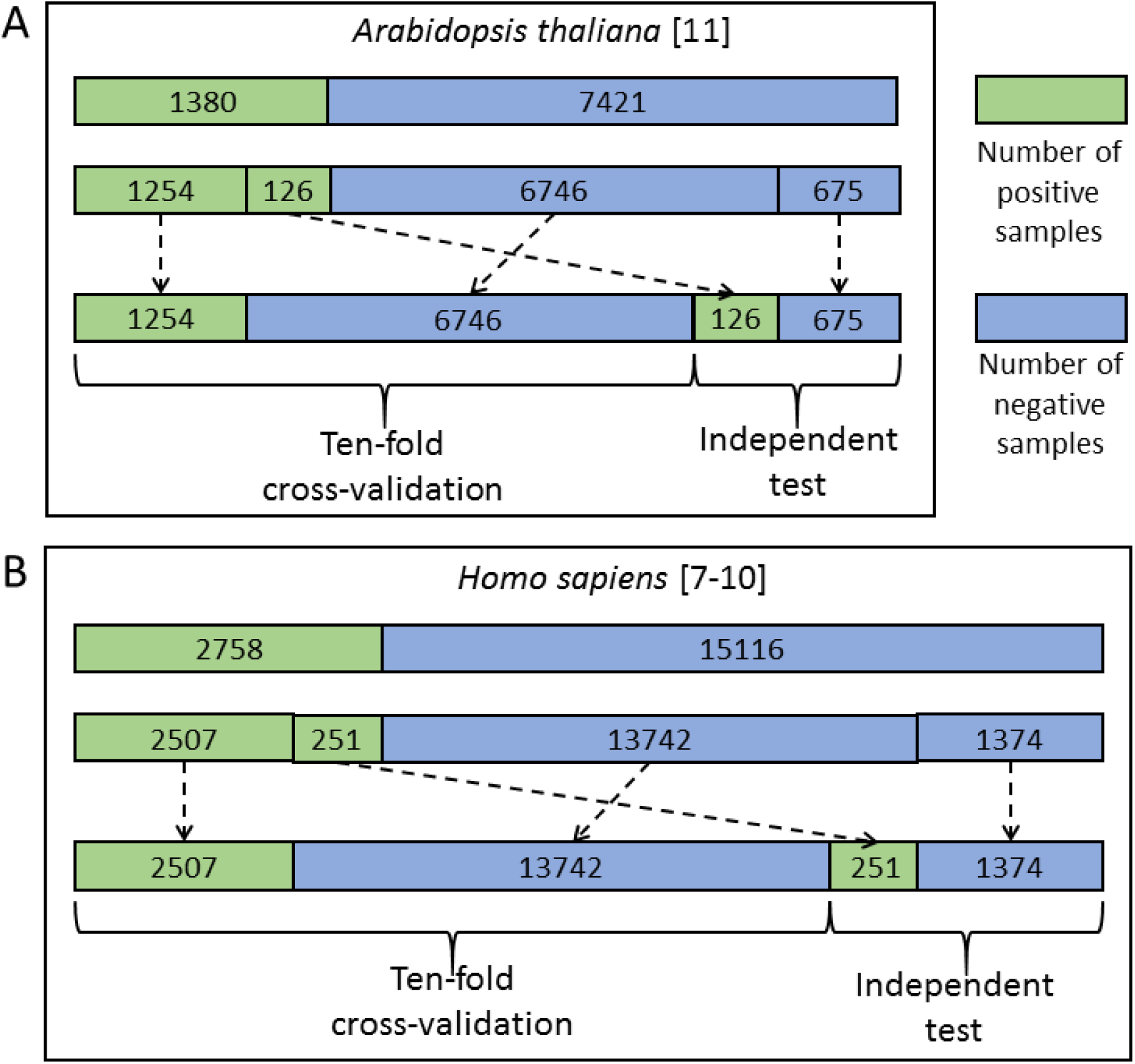
The flowchart of dataset process for *A. thaliana* (A) and *H. sapiens* (B).

We mapped 1537 Arabidopsis CSO sites [13] to the UniprotKB database[30] and 1535 sites from 1,130 proteins were retained as positive sites. The rest 8819 Cysteine residues in the same proteins were defined as negative sites. Moreover, we truncated these protein sequences into 35-residue segments with the Cysteine located at the center and the positive/negative sites correspond to positive/negative segments, respectively. It should be noted that if the central Cysteine was located around the N-terminus or C-terminus of a protein sequence, the gap symbol ‘-’ was added to the corresponding positions to ensure that the segment had the same window size. The window size was optimized as a hyper-parameter in the Bayesian optimization method (Table S2) and finally determined as 33. Furthermore, to reduce the potential influence of the segments with high similarity on the performance of the models to be constructed, we set the identity of any two sequences with less than 40%, referring to previous studies [14, 15, 22]. When the identity was >40% between two positive segments or two negative segments, one was randomly removed. When the identity was >40% between a positive and a negative segments, the positive was retained and the negative was discarded. As a result, 1380 positives and 7421 negatives were retained. Finally, we randomly separated the positive and negative segments into 11 groups of which ten were used for ten-fold cross-validation (1254 positives and 6746 negatives) and the rest for independent test (126 positives and 675 negatives) (Figure 1A). Similarly, the cross-validation dataset for *H. sapiens* contained 16249 samples (2507 positives and 13742 negatives) and the independent test set comprised 1625 samples (251 positives and 1374 negatives) (Figure 1B). These datasets are available at http://www.bioinfogo.org/DeepCSO/download.php.

### 2.2 Feature encoding schemes

#### Numerical representation for amino acids (NUM)

The NUM encoding approach maps each type of amino acid residues to an integer [31]. Specifically, in the alphabet “AVLIFWMPGSTCYNQHKRDE-”, each letter from ‘A’ to ‘-’ is converted to the integers from 0 to 20 in turn. For example, the sequence “VAMR” is encoded as “1,0,6,17”.

#### Enhanced Amino Acid Composition (EAAC)

The EAAC encoding [26, 32–34] introduces a fixed-length sliding window based on the encoding of Amino Acid Composition (AAC), which calculates the frequency of each type of amino acids in a protein or peptide sequence [35]. EAAC is calculated by continuously sliding a fixed-length sequence window (using the default value as 5) from the N-terminus to the C-terminus of each peptide. The related formula is listed below:

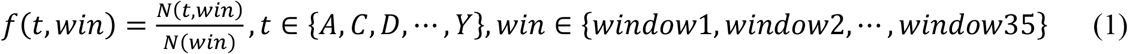

where *N(t, win)* is the number of amino acid *t* in the sliding window *win*, and *N(win)* is the size of the sliding window *win*.

#### Binary encoding

In the binary encoding, each amino acid is represented by a 21-dimensional binary vector [36]. The 21 dimensions are represented 20 amino acids and a complement “-” separately. The corresponding position is set as 1 and the rest position is set as 0. For example, the amino acid “A” is represented by “100000000000000000000”, “V” is represented by “010000000000000000000”, and the symbol “-” is represented by “000000000000000000001”, according to the alphabet “AVLIFWMPGSTCYNQHKRDE-”.

### 2.3 Optimization method for hyper-parameters

The hyper-parameters of a ML classifier affect the prediction performance. However, there are no formal rules to find the optimal hyper-parameters. Therefore, a lot of combinations of hyper-parameters need to be tested, which is time-consuming and tedious. We developed a simple method to automatically adjust and evaluate hyper-parameters (Figure 2A). This method contained two search approaches: grid search and Bayesian Optimization (BO).

**Figure 2.**
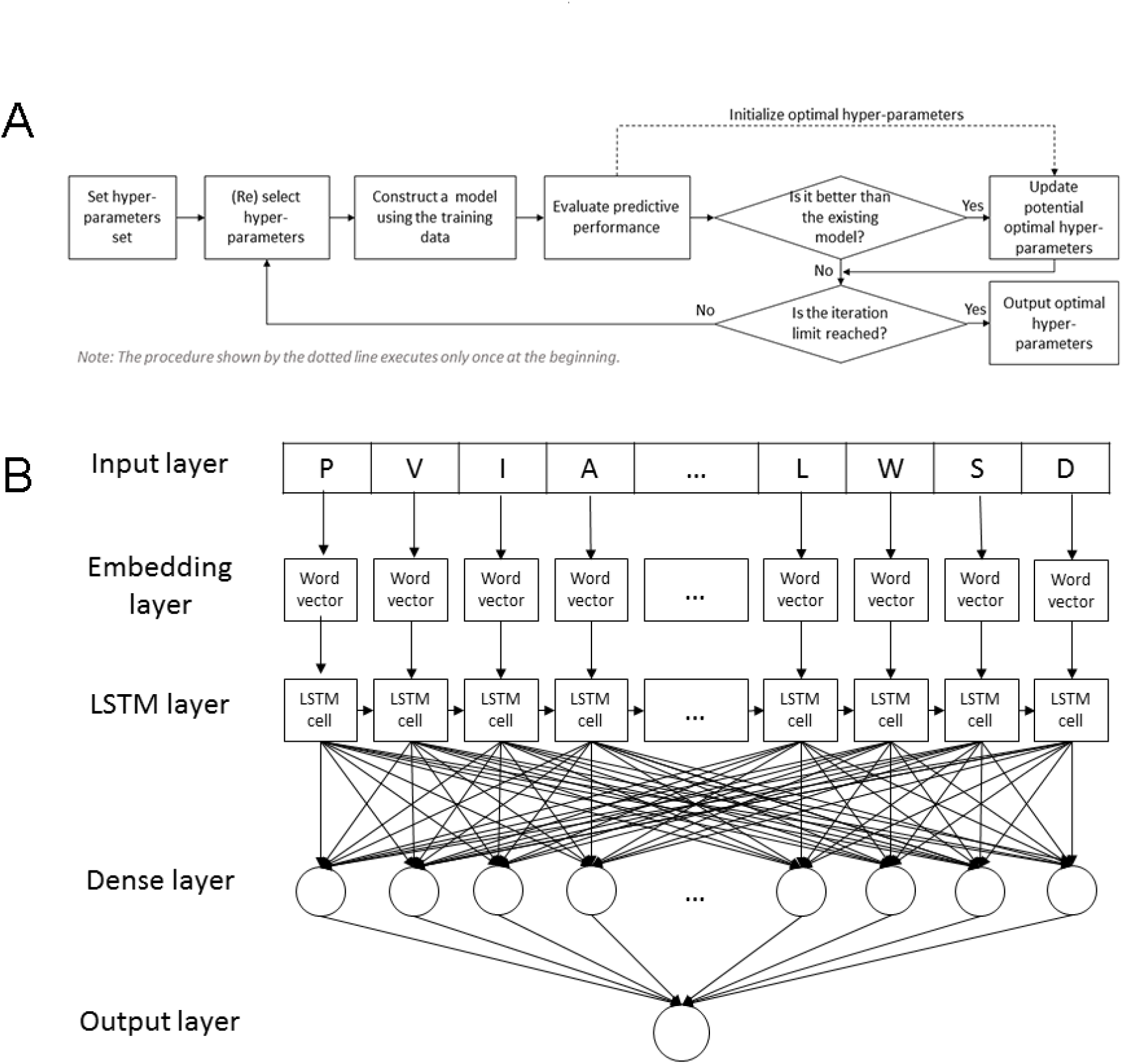
Hyper-parameter optimization for machine-learning classifiers (A) and the LSTM_WE_ architecture (B).

Grid search is a brute-force method to find the optimal hyper-parameters by training models using each possible combination of hyper-parameters and retaining the hyper-parameters corresponding to the model with the best performance. The grid search method is applicable to limited number of hyper-parameters due to the exponential increase in time spent with the number of hyper-parameters. In this study, Grid search was used for the RF-based models and the setting of hyper-parameter space was shown in Table S2. On the contrary, Bayesian Optimization (BO) provides a principled technique based on Bayes Theorem to direct a search of a global optimization problem that is efficient and effective. It is often used in deep learning to tune the hyper-parameters. The BO-related hyper-parameter space was listed in Table S2.

### 2.4 Architecture of the machine-learning models

#### 2.4.1 The LSTM model with the word embedding encoding approach (LSTM_WE_)

LSTM_WE_ contained five layers as follows (Figure 2B).

1. Input layer. Each peptide segment is converted into an integer vector with the NUM encoding.
2. Word Embedding layer. Each integer of the vector from the input layer is encoded into a four-dimension word vector.
3. LSTM layer. Each of the word vector is input sequentially into the LSTM cell that contained 32 hidden neuron units.
4. Dense layer. It contains a single dense sublayer that has 32 neurons with the ReLU activation function.
5. Output layer. This layer has only one neuron activated by sigmoid function, outputting the probability of the CSO modification.

#### 2.4.2 The CNN model with the word embedding layer (CNN_WE_)

CNN_WE_ (Figure S1), contains five layers, where the first two layers and last one layer are as same as LSTM_WE_. The third layer is 1D convolution layer with 20 filters and kernel size as nine and the fourth layer has a single dense sublayer with 16 neurons. The optimal hyper-parameter values are obtained using Bayesian optimization algorithm.

#### 2.4.3 The RF algorithms with different features

The RF algorithm integrates multiple decision trees and chooses the classification with the most votes from the trees. Each tree depends on the values of a random vector sampled independently with the same distribution for all trees in the forest. In this study, we constructed the RF models with three different features, including binary encoding, EAAC encoding and word embedding. The number of decision trees was selected as 580 *via* the grid search method. These classifiers were developed based on the Python module “sklearn”.

### 2.5 Performance assessment of the predictors

To evaluate the performance of our proposed method, four measures including accuracy (ACC), specificity (SP), sensitivity (SN) and the Matthew’s correlation coefficient (MCC) were used. They are defined as follows:

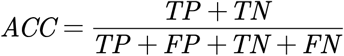

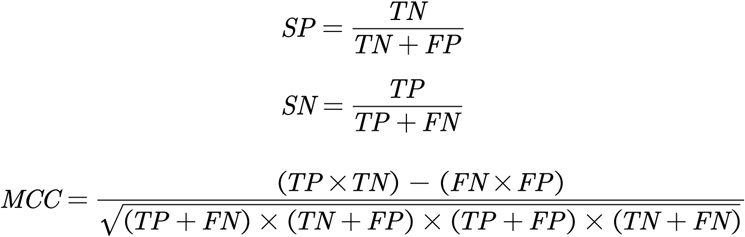

Where TP, TN, FP and FN represent the true positives, true negatives, false positives and false negatives respectively. Additionally, due to the number of positive and negative samples used in the experiment was unbalanced, and the above indexes were calculated based on the threshold value. In order to evaluate our model more objectively, an index independent of the threshold value and not affected by the sample ratio was needed. Therefore, the receiver operating characteristic (ROC) curves, together with the AUC are employed to comprehensively evaluate classification performance. Specifically, due to low false positive rate of a predictor is significant in practical application, the area under ROC curve with <10% false positive rate (AUC01) was considered.

### 2.6 Statistical methods

The paired student’s t-test was used to test the significant difference between the mean values of the two paired populations. The adjusted P value with the Benjamini-Hochberg (BH) method was adopted for multiple comparisons.

## 3 Results and discussion

Several computational approaches has been developed for the prediction of human CSO sites [37, 38]. Recently, this modification has been investigated across different species and the number of reported sites has been significantly increased. These raised our interest to develop novel prediction algorithms and explore the characteristics of this modification. The major procedure of our analysis contained three steps (Figure 3): data collection and preprocessing, in which the sample data were separated into the cross-validation dataset and the independent test dataset for model construction and evaluation; classifier construction, which involves data decoding, model training and hyper-parameter adjustments for resulting in a robust predictive model; the development of the final model as an online prediction tool.

**Figure 3.**
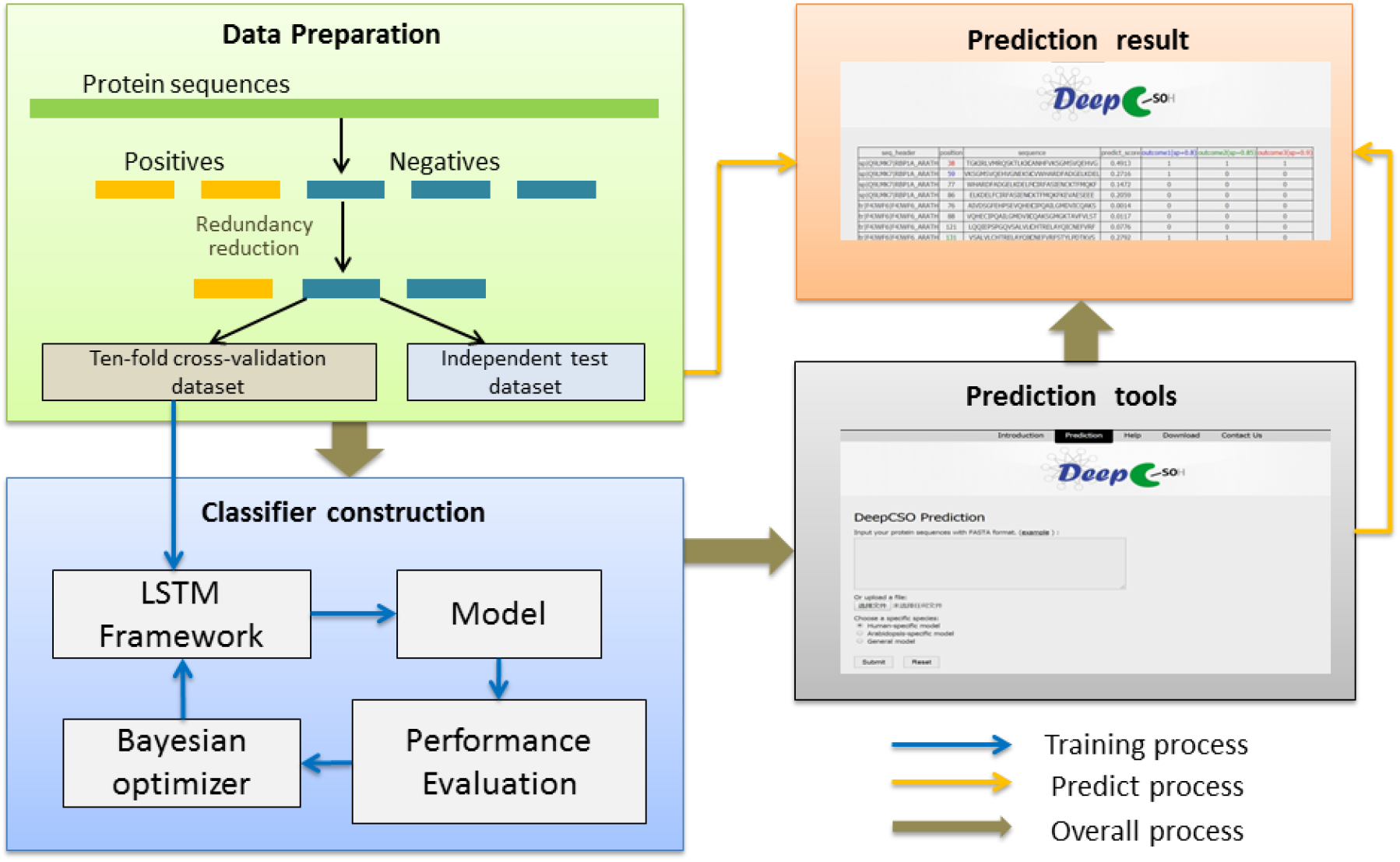
The flowchart of the prediction model construction.

### 3.1 LSTM_WE_ classifier performed favorably to other ML classifiers

Many computational approaches for predicting PTM sites are generally based on traditional ML algorithms combined with various features encoded from peptide sequences. The RF model is widely applied due to its robustness to data imbalance[39]. Accordingly, we constructed RF-based predictors with different encoding schemes for the CSO prediction. The encoding schemes include binary and EAAC and the corresponding classifiers were dubbed RF_BINARY_ and RF_EAAC_. Moreover, deep learning (DL) algorithms have recently been applied to the field of the modification prediction and demonstrated their superior performances [33, 40]. Accordingly, we developed two different DL classifiers based on word embedding [41], named as CNN_WE_ and LSTM_WE_.

We first took the *Arabidopsis* data to construct and compare different models [13]. The *Arabidopsis* cross-validation dataset contained 8000 samples (1254 positives and 6746 negatives) and the independent test set covered 801 samples (126 positives and 675 negatives) (See methods for details; Figure 1). We compared the performances of these algorithms in terms of Acc, Sn, MCC, AUC and AUC01 values for both the ten-fold cross validation and the independent test (Table 1). In our previous studies DL models showed superior performance than traditional ML models [33, 42]. It is still true for the CSO prediction. LSTM_WE_ had the best performance among these constructed models in terms of Acc, Sn, MCC and AUC values for both ten-fold cross validation and independent test. For instance, Its AUC value is 0.852 for the cross-validation and its values of Acc, Sn, Sp and MCC were 0.786, 0.717, 0.799 and 0.417, respectively (Table 1, Figure 4A&C). As prediction performance at a low false positive rate is highly useful in practice, we estimated these predictors using AUC01, where the specificity was determined to be >90%. LSTM_WE_ again showed the largest AUC01 values for both ten-fold cross-validation and the independent test (Figure 4B&D). As the encoding approach has great impact to the traditional ML models [33, 42, 43] and the WE approach integrated with LSTM had the best performance in this study, we attempted to investigate whether the integration of WE and RF had a good performance. Accordingly, we extracted embedding layer vector as feature encoding from LSTM_WE_ and trained the RF model, dubbed RF_WE_. Interestingly, RF_WE_ did not show good performance compared to RF_EAAC_, CNN_WE_ or LSTM_WE_. It suggests that the WE encoding approach may be improper for the construction of traditional ML algorithms.

**Table 1.**
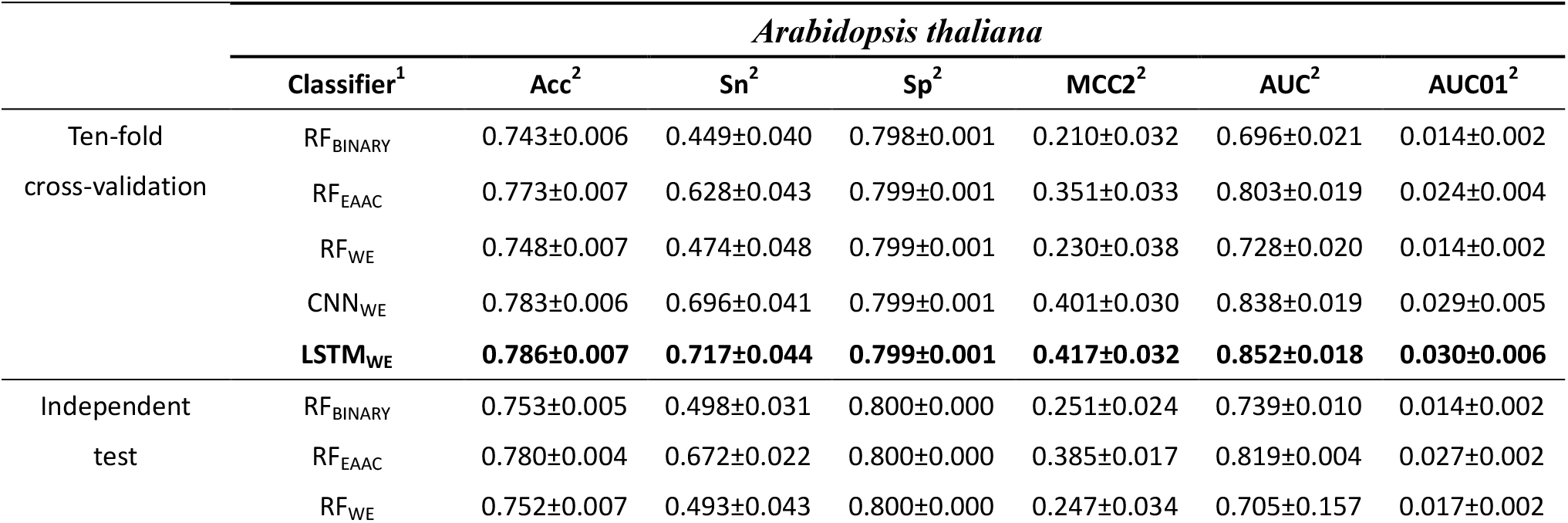
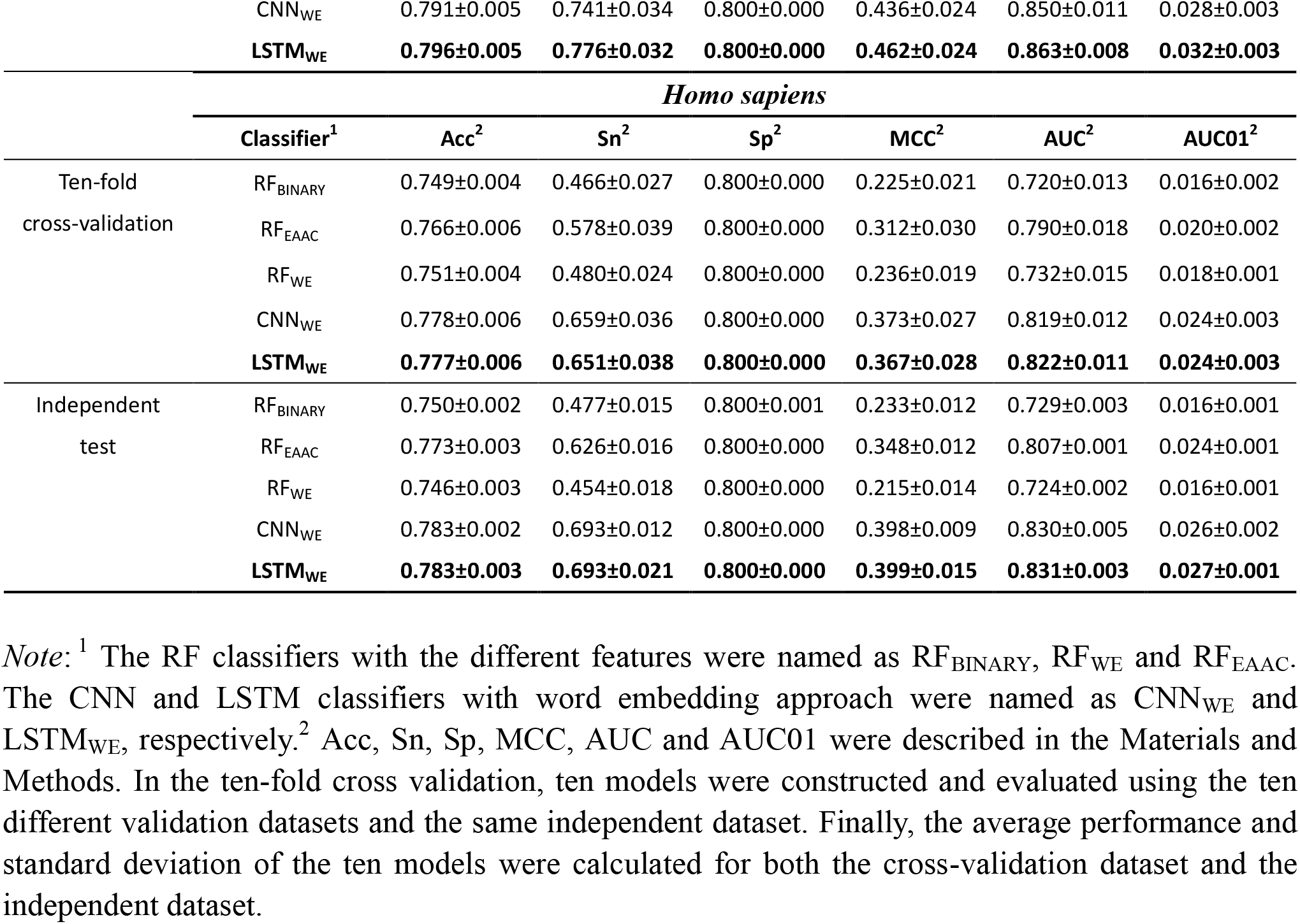
Performances of the different classifiers for *Arabidopsis thaliana* and *Homo sapiens*

**Figure 4.**
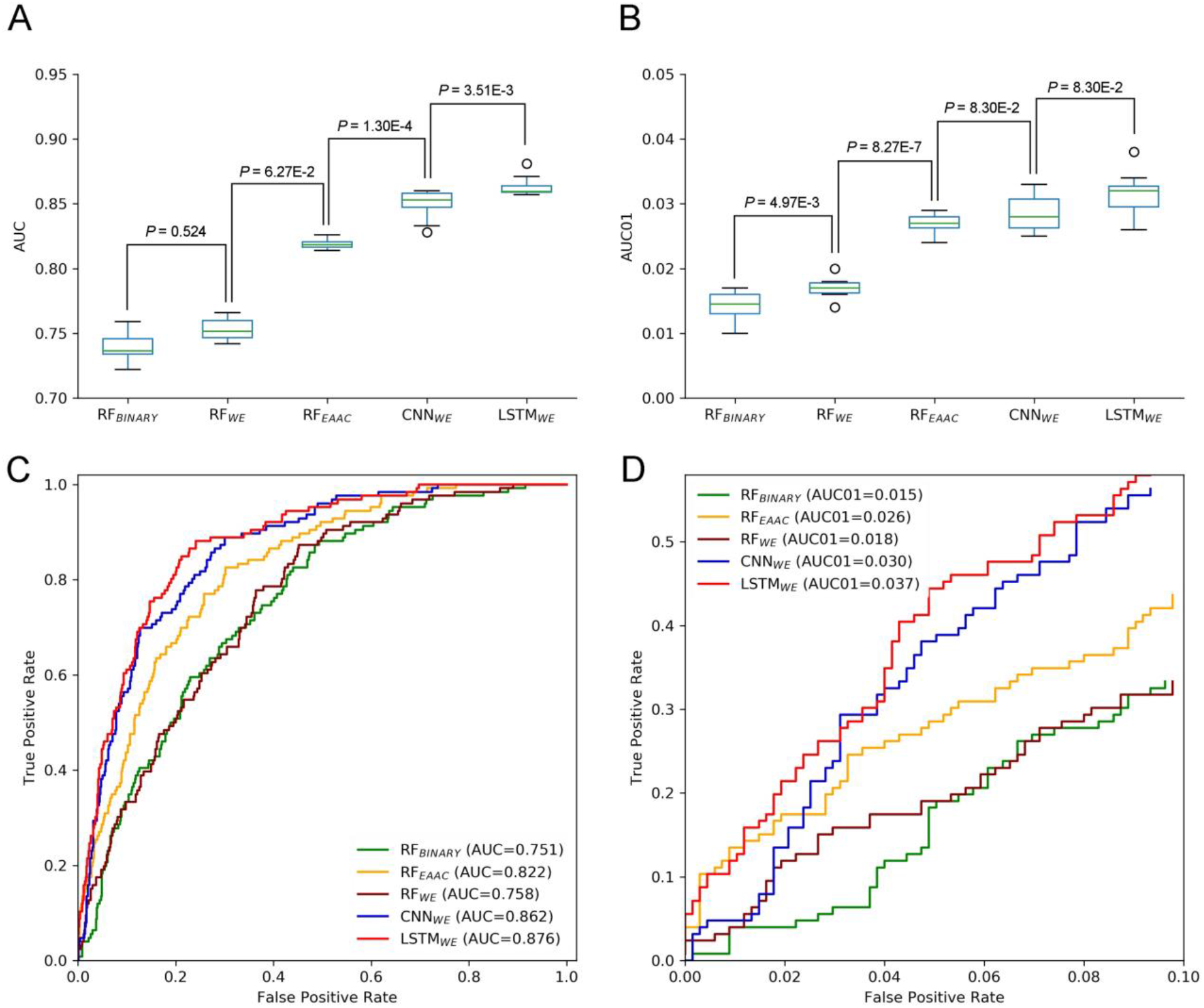
Performance comparison of different CSO predictors on *Arabidopsis thaliana*. The performances of CSO predictors were compared in terms of AUC (A) and AUC01 (B), respectively, for ten-fold cross-validation. AUC (C) and AUC01 (D) curves were generated using the independent test.

We further constructed the models for the human organism. The Human cross-validation dataset contained 16249 samples (2507 positives and 13742 negatives) and the independent test set covered 1625 samples (251 positives and 1374 negatives) (Figure 1B). Similarly, LSTM_WE_ had the best performance (Table 1, Figure S2). For instance, its values of AUC, Acc, Sn, Sp, MCC and AUC01 for the cross-validation were 0.822, 0.777, 0.651, 0.800, 0.367 and 0.024, respectively. We evaluated the robustness of LSTM_WE_ by comparing their performances between the cross-validation and independent tests for individual organisms. As their performances between these two tests had no statistically different for each species (P=0.18/0.085 for the arabidopsis/humans, respectively), we concluded that our constructed models were robust and neither over-fitting nor under-fitting (Table 1).

### 3.3 Human-specific LSTM_WE_ performed better than reported classifiers

Several computational methods have been reported for the prediction of human CSO sites (See Introduction for details). We compared these methods with LSTM_WE_ using the cross-validation results derived from the original studies, which were developed based on the CSO sites identified using the chemoproteomics approach (Table 2). LSTM_WE_ also showed the best performance in terms of ACC, MCC and AUC (Table 2). As iSulf-Cys[15] is the only accessible model to date, we compared it and LSTM_WE_ using the human independent dataset of this study. The AUC value (0.839) of LSTM_WE_ is significantly larger than that (0.666) of iSulf-Cys (Figure S3). In summary, LSTM_WE_ performed better than the reported classifiers.

**Table 2.** The k-fold cross-validation results of existed tools

| Tools | k-fold | Acc | Sn | Sp | MCC | AUC |
| --- | --- | --- | --- | --- | --- | --- |
| **DeepCSO** | **10** | **0.777±0.006** | **0.651±0.038** | **0.800±0.000** | **0.367±0.028** | **0.822±0.011** |
| MDD – SOH | 5 | 0.68 | 0.7 | 0.7 | 0.27 |  |
| SOHSite | 5 | 0.71 | 0.72 | 0.72 | 0.3 |  |
| SOHPRED | 5 |  | 0.727±0.005 | 0.742±0.001 | 0.336±0.004 | 0.801±0.001 |
| iSulf-Cys | 10 | 0.656±0.007 | 0.673±0.007 | 0.639±0.001 | 0.312±0.014 | 0.716±0.009 |
| SulCysSite | 10 |  | 0.745±0.006 | 0.744±0.002 | 0.343±0.005 | 0.806±0.002 |
| Sulf_FSVM | 10 | 0.711±0.002 | 0.733±0.004 | 0.708±0.002 | 0.297±0.004 | 0.788±0.002 |

### 3.4 Conservation of the CSO modification and the development of general LSTM_WE_ models

Cysteine S-sulphenylation has been identified across various organisms, ranging from yeasts to worms and from plants to humans [7, 8]. To understand its conservation, we compared the characteristics of CSO-containing peptides in human and arabidopsis species, respectively, using the two-sample-logo approach [44]. Figure 5 showed that both species shared the enriched basic amino acids R and K and the depleted polar neutral amino acid C. Nevertheless, the amino acid H is enriched for *A. thaliana* whereas the hydrophobic amino acid L was depleted for *H. sapiens*. As the characteristics of CSO-containing peptides are similar between both species, we hypothesized the generalization ability of our developed models. To test this hypothesis, we used the human LSTM_WE_ model to predict the arabidopsis independent test dataset and employed the Arabidopsis LSTM_WE_ model to predict the human independent test dataset. The AUC values were 0.799 and 0.766, respectively, significantly larger than the random prediction (*ie.* AUC=0.5; Table 3). Nevertheless, the cross-species prediction had relative low performance compared to the self-species prediction (AUC=0.876/0.839 for arabidopsis/human, respectively). As the CSO sites have been systematically analyzed in a few species, it is necessary to develop a general CSO prediction model according to its conservation to boost the investigation for other species. Accordingly, we mixed the training datasets of *H. Sapiens* and *A. thaliana* and constructed the general LSTM_WE_ model and validated it using the independent datasets from both organisms. The performance of the general LSTM_WE_ model is slightly lower than that of the self-species prediction, which may be caused by the interference of the CSO characteristics of other species (Table 3). Overall, the conservation of the CSO modification leads to the effective prediction of the general LSTM_WE_ classifier.

**Table 3.**
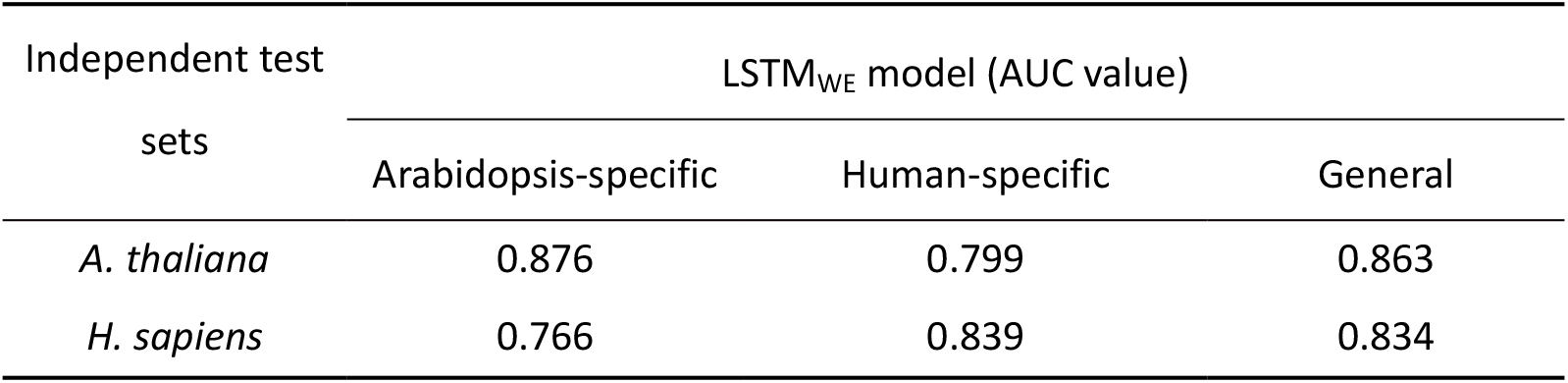
Evaluation of species-specific and general LSTM_WE_ models using the independent test sets from different species.

**Figure 5.**
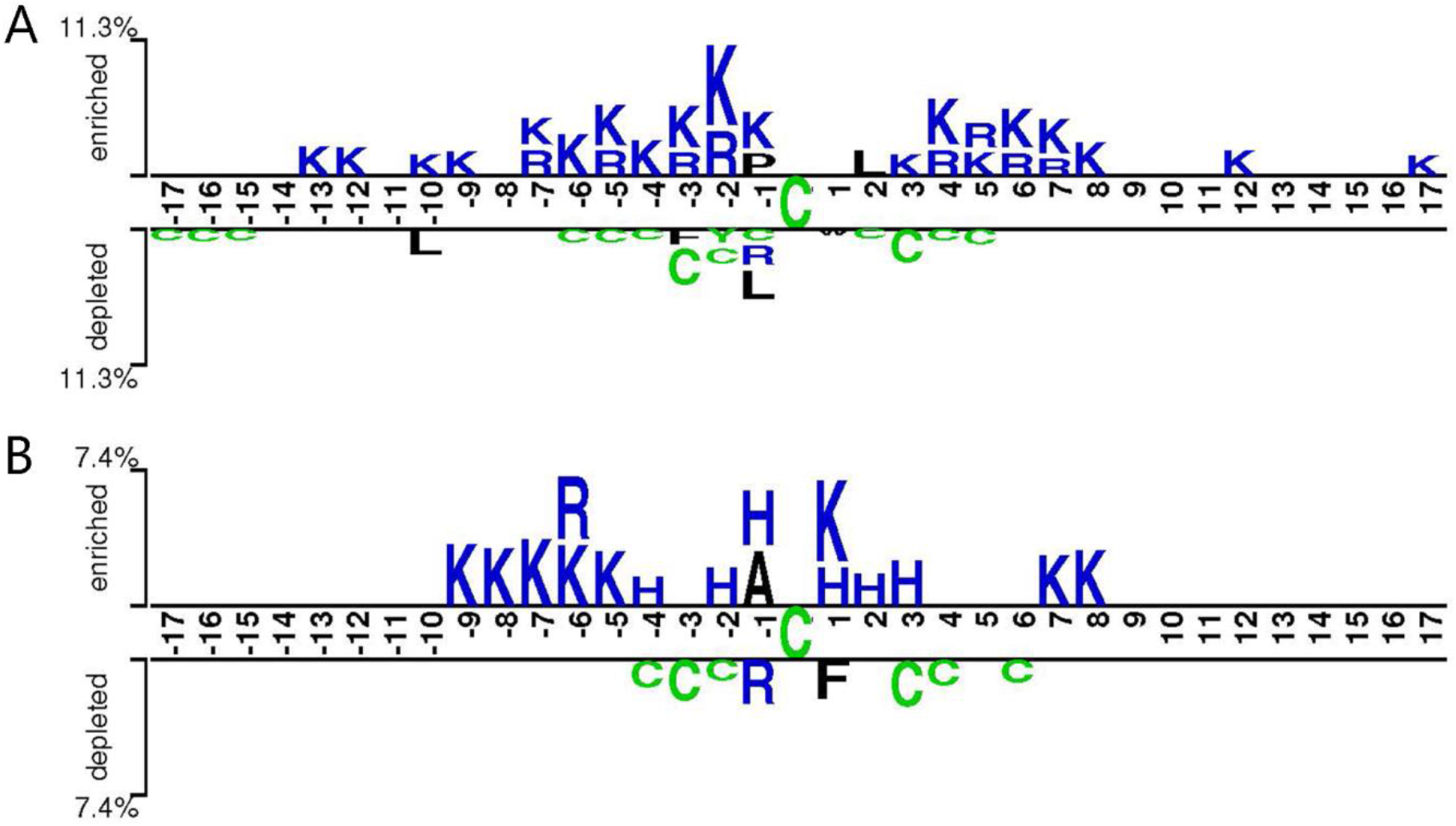
Sequence pattern surrounding the CSO sites, including the significantly enriched and depleted residues based on CSO-containing peptides and non-modification peptides for *H. sapiens* (A) and *A. thaliana* (B) (P<0.05, t-test with Bonferroni correction). The pattern was generated using the two-sample-logo method ([44]).

To further understand the performance of the general LSTM_WE_ classifier, we visualized the sample distributions, based on the human independent dataset, from the outputs of the input layer, embedding layer, LSTM layer and dense layer of the general model using t-SNE algorithm[45] (**Figure 6**). After the input layer (**Figure 6A**), the positive and negative samples were mixed together, as the training goes on (**Figure 6B&C**), positive and negative samples were gradually separated. After the LSTM layer, they were clearly separated (**Figure 6D**). This comparison indicates that the LSTM layer is a powerful method to detect the distinctive features of the positives and negatives. The similar observation is made for the arabidopsis independent test dataset (Figure S4).

**Figure 6.**
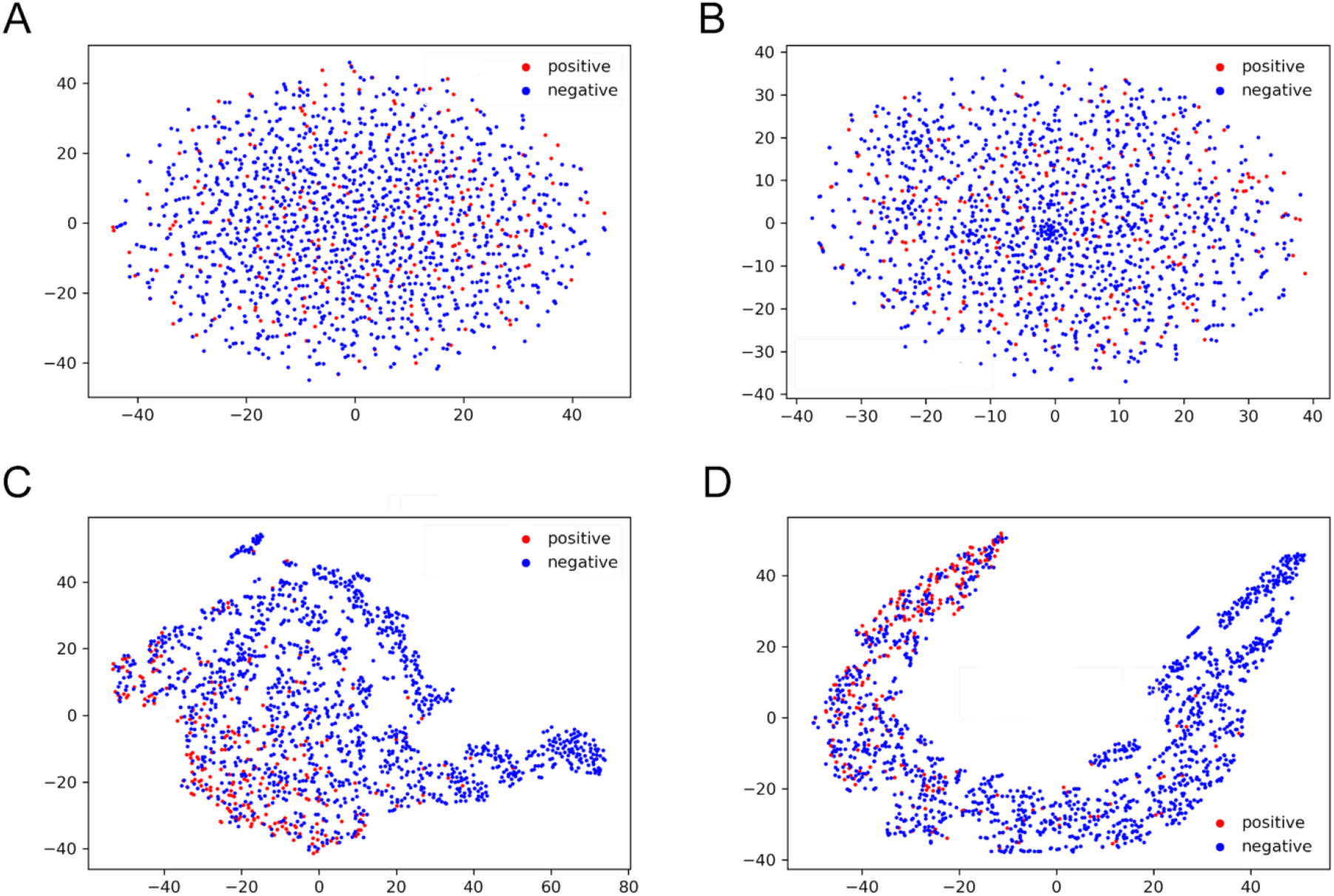
T-SNE visualization of the distributions of peptides in the human independent dataset for the outputs of input layer (A), embedding layer (B), LSTM layer (C) and dense layer 2 (D) of the general LSTM_WE_ model.

### 3.4 Construction of the on-line CSO predictor

We developed an easy-to-use online tool for the prediction of the CSO sites, dubbed DeepCSO. DeepCSO contains three LSTM_WE_ models: the general model and two species-specific models (*ie. H. sapiens* and *A. thaliana*). The users could select the general model or species-specific model at the input interface and input the query protein sequences directly or upload the sequence file. After the job submission, the prediction will start and the prediction process may take several minutes. Finally, the prediction results are output in tabular form with five columns: sequence header, position, sequence, prediction score, and prediction results at the specificity levels of 80%, 85% and 90%, respectively.

## 4 Conclusions

The currently CSO predicted tools are based on traditional machine learning method that requires experts to pre-define informative features, and there was no prediction tool have been developed for other than human species. We developed three LSTM-based prediction models, where two are organism-specific and one is general, and they compared favorably to the reported models. Despite lacking pre-defined features, the deep learning classifier demonstrated a superior performance compared to the traditional machine learning methods. This may be due to the self-learning ability of deep learning. The outstanding performance of our general model suggests that the Cysteine S-sulphenylation is well conserved and the LSTM-based model has an advantage in long-term memory to capture the key features of the entire sequences.

---

**Table S1.** Summary of the experientially identified CSO sites reported in literature

| Species | Cell lines | Number of CSO sites | PMID |
| --- | --- | --- | --- |
| *Homo sapiens* | HeLa | 1098 | 30177848 |
| *Homo sapiens* | A549 | 1173 | 30177848 |
| *Homo sapiens* | RKO | 1105 | 25175731 |
| *Homo sapiens* | RKO | 1283 | 28355876 |
| *Homo sapiens* | THP-1 | 268 | 27690452 |
| *Arabidopsis thaliana* |  | 1537 | 31578252 |

**Table S2.** Hyper-parameters optimization scheme

| **Grid search space** |
| --- |
| `param_test= { 'n_estimators':range(100,1001,20), 'max_depth':range(6,15,2), 'min_samples_split':range(2,9,1) }` |
| *Note: the above range was divided into three sections according to the last test results.*<br>`gsearch= GridSearchCV( estimator = RandomForestClassifier(oob_score=True, random_state=10), param_grid =param_test, n_jobs=-1, scoring='roc_auc', iid=False, cv=3, verbose=2, refit=True )` |
| **Bayesian optimization space** |
| `space = { 'window': hp.choice('window', [35,33,31,29,27,25,23]), 'emb': hp.choice('emb', [3,4,5,6,7]), 'lstm': hp.choice('lstm', [8,16,32,64]), 'filters': hp.choice('filters', [18,20,22,24]), 'kernel_size':hp.choice('kernel_size',[8,9,10,11,12]), 'units': hp.choice('units', [8,16,32,64]), 'batch_size': hp.choice('batch_size', [16,32,64,128]), 'lr_decay': hp.loguniform('lr_decay', np.log(0.000001), np.log(0.001)), 'dropout_rate': hp.uniform('dropout_rate', 0.0, 0.5), 'epochs': hp.choice('epochs', [50,60,70,80,90,100,110,120,130,140,150]), }` |

**Figure S1.**
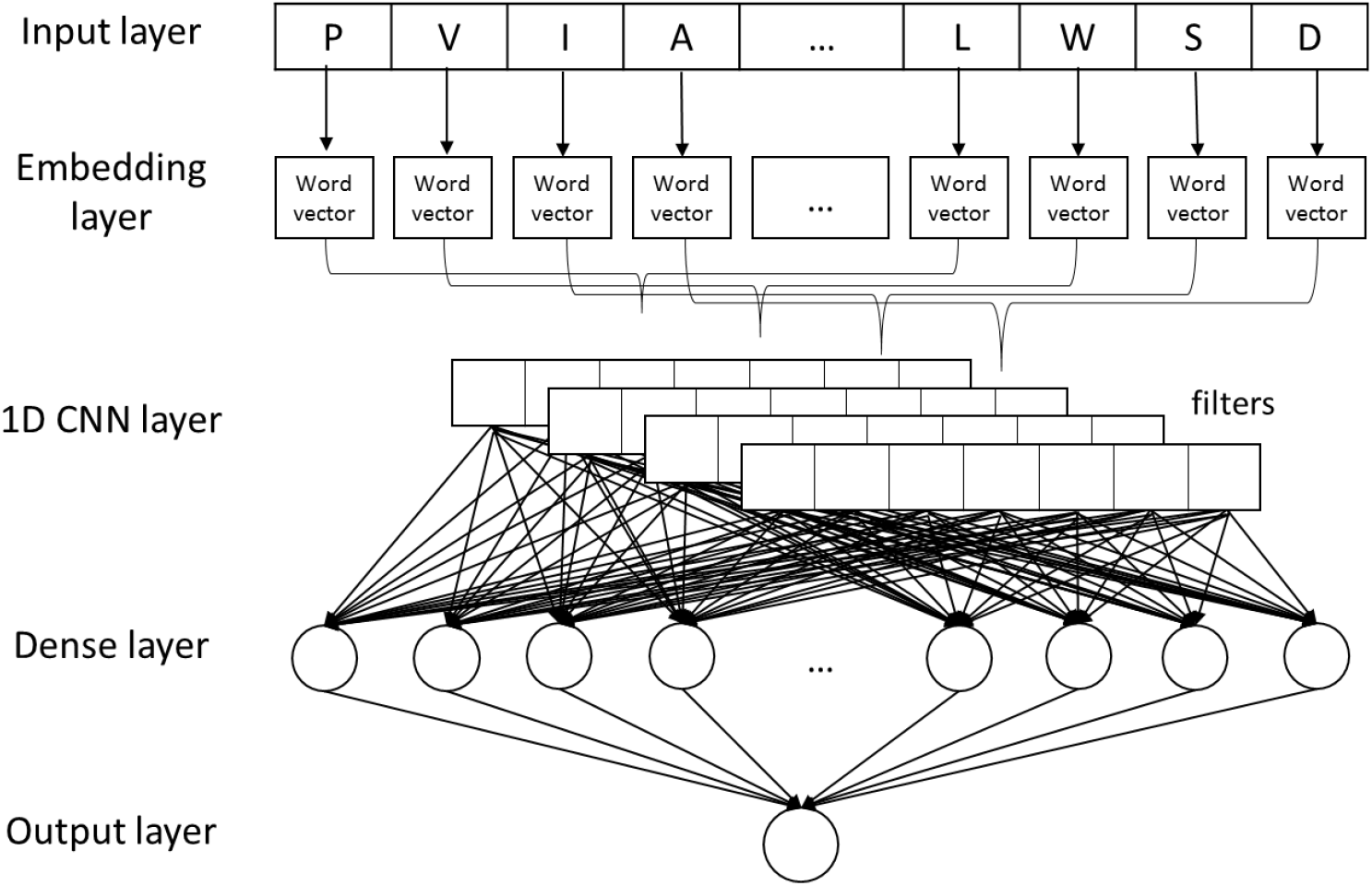
The CNN_WE_ architecture.

**Figure S2.**
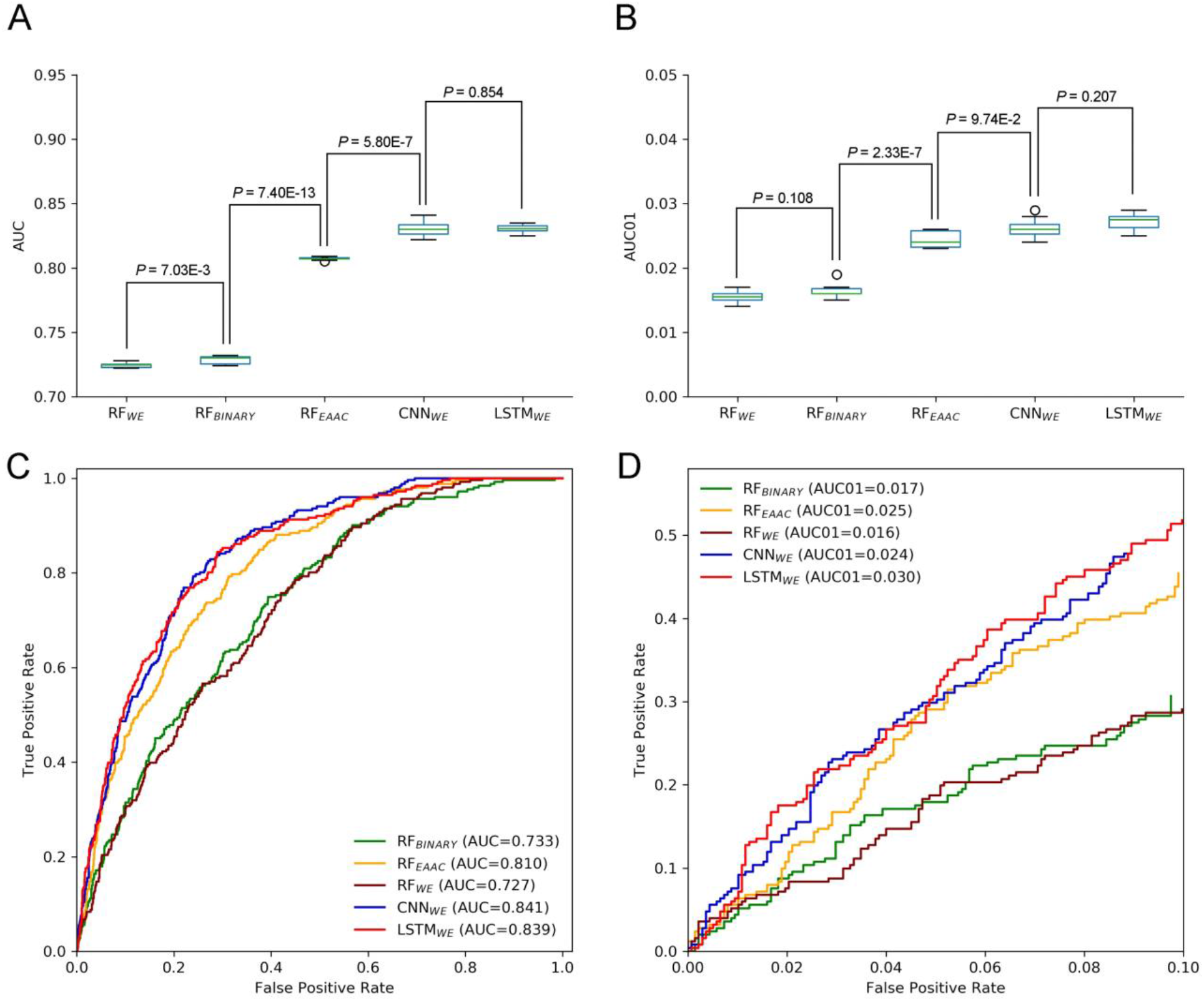
Performance comparison of different CSO predictors on *Homo sapiens*. The performances of CSO predictors were compared in terms of AUC (A) and AUC01 (B), respectively, for ten-fold cross-validation. AUC (C) and AUC01 (D) curves were generated using the independent test.

**Figure S3.**
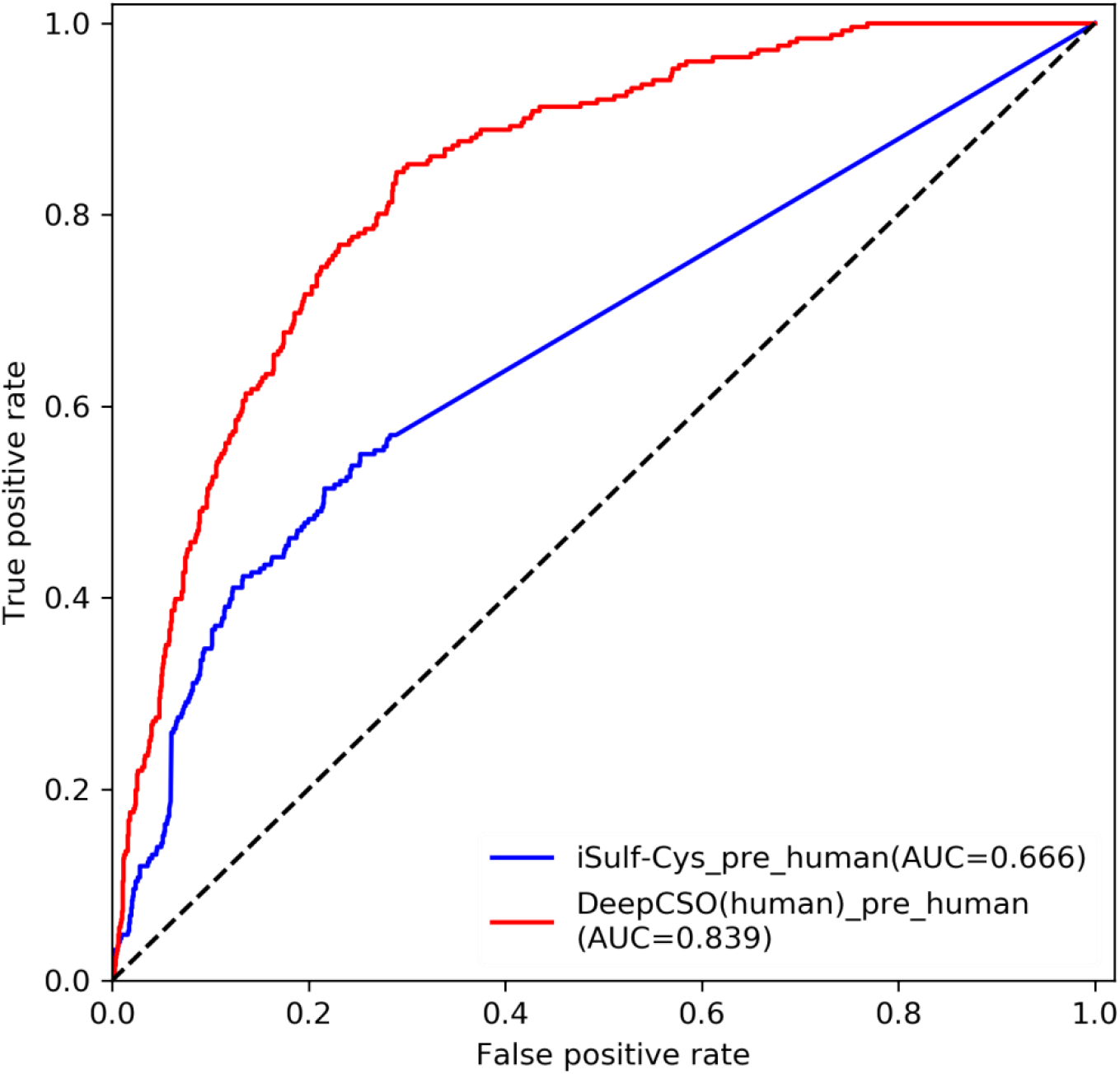
Performance comparison between LSTM_WE_ and iSulf-Cys in terms of the human independent dataset.

## Notes

http://www.bioinfogo.org/DeepCSO

### Competing Interest Statement

The authors have declared no competing interest.

